# Diallel Analysis for Morphological and Biochemical Traits in Tomato Cultivated Under the Influence of Tomato Leaf Curl Virus

**DOI:** 10.1101/371013

**Authors:** Prashant Kaushik, Major Singh Dhaliwal

## Abstract

Eloquent information about the genetic basis of inheritance is important for any breeding program. Therefore, a diallel study was conducted under the influence of tomato leaf curl virus (TLCV) disease, using the eleven advanced lines of tomato. Firstly, the information regarding percent disease index (PDI) was determined via artificial screening with viruliferous whiteflies. Later, these lines were crossed in half diallel mating design to produce fifty-five one-way hybrids. These hybrids and parental genotypes were evaluated for seven morphological and three biochemical traits under open field conditions. Using the Griffing approach (Method II and Model I) basis of inheritance of traits were determined. Also, a Bayesian model was applied to the total yield descriptor. Correlations data indicated that total yield was not correlated with any other trait. The significant general combining ability (GCA) and specific combining ability (SCA) values indicates exploitable genetic variation. The broad-sense heritability values were larger than narrow-sense heritability, showing that selection will be efficient for the improvement of these traits. Hybrid combinations H23, H42 and H49 can be considered efficient for the selection of multiple traits, including yield. Overall, this study provides a useful information regarding the genetics of important traits of tomato under TLCV infestation.

## 1. Introduction

Tomato (*Solanum lycopersicum* L.) is among the most cultivated plants; hence, the efforts of its genetic improvement dates back to last century and are still enduring through traditional breeding and genomics-based approaches [1,2]. Tomatoes are well acclimatized and breed to yield under extreme climatic conditions like drought and frost [3–5]. But, insect pest and diseases are still big challenges for the successful production of tomatoes. Worldwide, approximately around 146 viruses belonging to 33 different genera are reported to infect tomato plant [6,7]. Among them, the genus *Begomovirus* causes huge economic losses to tomato production. Belonging to this genus a DNA virus known as the tomato leaf curl virus (TLCV) is a serious disease of tomato and its incidence, can easily result up to 90 percent yield loss to the tomato crop [8,9].

Tomato production especially of autumn season crop in Northern India and summer season crop of Southern India is susceptible to the high incidence of tomato leaf curl virus (TLCV) disease [10]. TLCV is transmitted by whitefly(*Bemisia tabaci* Genn.) in a circulative and persistent manner [11]. Hitherto, in North India, tomato leaf curl New Delhi virus a strain of TLCV reported from New Delhi region of India has an extensive distribution and also infects several other vegetable crops e.g. Eggplant, Squash and Pepper [12]. As a response to TLCV infection plant leaves shows symptoms like, curling of leaf margins, shrinking and thickening of leaf surface. While the overall plant become stunted in growth, with few and misshaped fruits [11]. Extensive efforts in the form of phenotypic screening has been taken in order to identify resistant genotypes. In this respect, the artificial cage inoculation using viruliferous whiteflies is the most competent and reliable method to carry out the screening for TLCV disease. Although with artificial screening, the plant gives a strong reaction response, than it might give under the field conditions[13,14].

Tomatoes are a important source of nutraceuticals like vitamins (C, K, B6 etc.), phenolic acids, and minerals (folate, manganese etc.). All of these are vital for human health and body development [15,16]. Particularly, its fruits contain one of the important dietary carotenoids known as lycopene, important for the prevention of chronic diseases like breast, lung and prostate cancer [17–19]. The lycopene content of tomato varies based on genotypes genetic makeup, cultivation environment, disease pressure, and genotype-by-environment interactions [20]. The nutraceutical properties of tomato fruit have industrialized the processing of tomatoes; commonly, tomatoes are processed as juice, ketchup, paste, and sauce [21]. These biochemical aspects of the tomato fruit have become an important goal of tomato breeding programs [22,23].

Hybrid development is a successful approach for vegetable improvement, especially for solanaceous vegetables. Also, to chalk out a breeding strategy for successful cultivation under TLCV infestation it is important to have information about the inheritance of traits under the prevailing conditions. Therefore, estimation general combining ability (GCA) and the specific combining ability (SCA) is important for genetic enhancement of the crop. But, the amount of variation in GCA and SCA values not totally rest on gene effects besides it also involves the gene structure of the parents involved [24]. Diallel matting design based on general linear model framed by Griffing [25] is a popular choice and widely accepted tool for identification of the hybrid combinations of interest in tomato and in other members of Solanaceae [26–28]. Previously studies indicated that leaf characteristics and foliar pubescence affect the feeding preferences of whiteflies [29,30]. Sometimes, Griffingˋs method is not adequate in case of missing data, imbalance, and outliers under the situations where chances of bias are too high to avoid. This is especially true with experiments carried out under disease pressure conditions [31,32].

The use of more rigorous Bayesian methods are used to overcome these limitations [33,34]. The BayesDiallel approach uses a Markov chain Monte Carlo (MCMC) sampling distribution which intern provides a greater flexibility, which further improves the biological interpretability of results. Therefore, in this study, the parameter of the total yield (kg/plant) was investigated with this Bayesian approach. Bayesian approach is not popular among the plant breeding community because of calculation limitations and complexity of statistics involved [35]. But, the different models based on Bayesian approach provides a more vigorous and detailed analysis of highly variable and complex trait like yield. An acquaintance of the genetics of important morphological traits under tomato leaf curl virus infestation conditions will be helpful for carrying out efficient selection and breeding.

Therefore, the objectives of this study were to determine GCA, SCA, and heritability of tomato genotypes crossed in a half diallel mating design. Further, the BayesDiallel approach was used to provide a more comprehensive analysis of diallel data generated for the total yield. By applying BayesDiallel approach first time on the total yield data of the diallel cross under TLCV conditions along with Griffing’s method, we aim to precisely estimate the combining ability and the heritability estimates and to suggest a robust approach aimed at tomato TLCV resistance breeding.

## 2. Materials and Methods

### 2.1. Plant Material and Artificial Screening

Eleven advanced lines of tomato developed at the Punjab Agricultural University (coordinates at 30°54′6.893″ N 75°48′27.989″ E), Ludhiana, India were artificially screened for resistance to TLCV disease during 2011-12. The artificial screening of seedlings (~ 2-3 week old) was carried out by challenging 25 plants of each of the eleven genotype with viruliferous whiteflies reared on the TLCV disease affected plant, the detailed method is provided elsewhere [26]. The disease reaction of genotypes was scored on the scale, where 0–10% resistant (R), >10–30% moderately resistant (MR), >30–70% susceptible (S), and >70–100% highly susceptible (HS) [36,37]. Further, the potential disease incidence (PDI) was measured as the (number of infected plants/total number of plants) × 100. While, the eleven genotypes were crossed in a diallel mating design during February-March of 2012 resulting in fifty-five one way F1 hybrids. Thereafter, the first cross combination, H1 is referred to as the first cross in the half diallel i.e. H1 (P1 × P2) and so on, until the last cross as H55 (P10×P11) (Table 1). The 11 parental genotypes and 55 one-way hybrids were evaluated under the whitefly infestation conditions during the August of 2012 in a randomized complete block design. Each entry was replicated twice and each replication accommodated 15 plants. Plant production practices were exercised as per the Package of Practices (Anonymous) and no chemical treatment was used to control the whiteflies. Data were recorded on 13 central plants leaving one plant on either side of the row.

**Table 1.** Representation of the cross or hybrid combinations (55 in total) developed as a result of half-diallel matting design.

|  | P1 | P2 | P3 | P4 | P5 | P6 | P7 | P8 | P9 | P10 | P11 |
| --- | --- | --- | --- | --- | --- | --- | --- | --- | --- | --- | --- |
| P1 |  | H1 | H2 | H3 | H4 | H5 | H6 | H7 | H8 | H9 | H10 |
| P2 |  |  | H11 | H12 | H13 | H14 | H15 | H16 | H17 | H18 | H19 |
| P3 |  |  |  | H20 | H21 | H22 | H23 | H24 | H25 | H26 | H27 |
| P4 |  |  |  |  | H28 | H29 | H30 | H31 | H32 | H33 | H34 |
| P5 |  |  |  |  |  | H35 | H36 | H37 | H38 | H39 | H40 |
| P6 |  |  |  |  |  |  | H41 | H42 | H43 | H44 | H45 |
| P7 |  |  |  |  |  |  |  | H46 | H47 | H48 | H49 |
| P8 |  |  |  |  |  |  |  |  | H50 | H51 | H52 |
| P9 |  |  |  |  |  |  |  |  |  | H53 | H54 |
| P10 |  |  |  |  |  |  |  |  |  |  | H55 |
| P11 |  |  |  |  |  |  |  |  |  |  |  |

### 2.2. Morphological and Biochemical Data Analysis

Data for seven morphological and three biochemical traits were recorded and inferences were made. Total yield (kg plant^−1^) and marketable yield (kg plant^−1^) were determined on per plant basis i.e. by diving the harvest of all picking with the number of plants. Fruit weight (g) was estimated as an average weight per fruit of the ten fruits sample collected at the red ripe stage. Five fruits per replication were used to measure the equatorial diameter, polar diameter and pericarp thickness. The polar diameter of cut fruits was measured as the distance between the stalk end and the blossom end. Conversely, the equatorial diameter was measured as the transversal distance of the fruit. Pericarp thickness was measured from centre of the fruit. These fruit-based morphometric measurements were recorded with the help of the Vernier caliper.

Three biochemical parameters estimated included dry matter, lycopene and total soluble solids (TSS) using three samples for each replication. Each sample was constituted of five red ripe fruits. Dry matter percentage was measured as the change of weight before and after oven drying at 70°C and was calculated based on the formula 100 × (dry weight / fresh weight). TSS content of fruits was determined with the help of a hand refractometer (RA-130-KEM, Kyoto Electronics Manufacturing Co., Ltd., Kyoto, Japan). The readings were recorded as °Brix (0 to 32) at room temperature. Lycopene content was determined by the method suggested elsewhere [38]. The optical densities of processed extracts for the lycopene content were recorded at 505 nanometers (nm) using Spectronic 20 (Thermo Fisher Scientific, Waltham, Massachusetts, USA). The combining abilities were estimated based on the Griffing’s Method 2 (parental genotypes and one-way hybrids) with Model 1 (fixed effect).[25] The diallel calculations were performed with the help of pacakge AGD-R (analysis of genetic designs with the R software package) version 4.0 [39]. Pearson coefficients of correlation (r) and there P-values were calculated and plotted using the packages corrplot[40]. The chart. Correlation() function within the PerformanceAnalytics package was used to generate scatter plots and histograms along with detailed information regarding correlations[41].

### 2.3. Bayesian Model-Based Analysis of Total Yield

The R package BayesDiallel was used for the Bayesian model-based analysis of total yield data [34]. Briefly we used the “Bab” model in the BayesDillel package, this model includes an overall effect of being inbred (B), an additive component (a), and a measure of parent-specific inbred deviation (b). Also, the “Bab” model was applied using an MCMC Gibbs sampler with five chains, 10,000 iterations, and burn-in of 1000 [35].

## 3. Results and Discussion

### 3.1. Artificial Screening

Artificial screening with viruliferous whiteflies provided a dissimilar percent of disease incidence (PDI) values for each genotype (Figure 1). The minimum PDI% was recorded in P4 (29.2%, MR), whereas the maximum was recorded with the P1 (100%, HS) (Figure 1). All the eleven parental genotypes exhibited the TLCV symptoms after 60 days of artificial inoculation with viruliferous whiteflies. On the TLCV disease scale, two lines showed the mild resistance (MR), in contrast, three were highly susceptible (HS), while the rest were susceptible (Figure 1).

**Figure 1.**
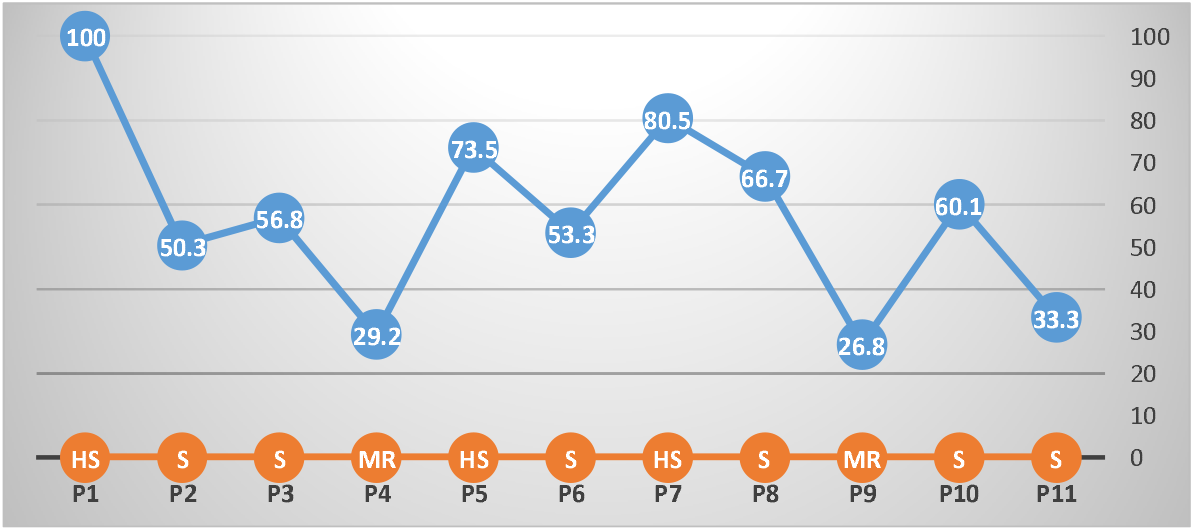
The percent disease incidence (PDI) reaction of parents to TLCV along with the grading scale (x-axis), where, HS (Highly Susceptible), MR (Moderately Resistant), and S (Susceptible).

Earlier studies have shown that as compared to a natural field environment screening of genotypes for TLCV, artificial screening is more useful as it ensures a uniform disease infection, and leave few chances for a susceptible plant to escape infection due to non-preference and loss of whiteflies infectivity. But, 100% disease incidence is also common with artificial inoculation method [13,42]. Artificial screening method using whiteflies is commonly applied to find out the reaction of tomato lines to TLCV disease [43–45]. Previously, the cultivated accessions and the landraces of tomato exhibited a varying range of the disease symptoms but lacked the complete resistance [46]. However, wild relatives of tomato e.g. *S. chmielewskii*, *S. habrochaites* and *S. pimpinellifolium* were found resistant even with artificial screening [14]. The differences in the results of field-based screening method could also be due to the difference in virus strain, vector genotype or altered feeding conditions [47, 48]

### 3.2. Variation in Parents and Hybrids

The magnitude of the mean squares of genotypes indicated that there were significant differences among the genotypes for all morphological characters studied pointing out the presence of genetic variability (Table 2). Similarly, significant GCA and SCA effects for all measured traits were detected (P ≤ 0.01). The lowest values of GCA/SCA ratio was noted for polar diameter and TSS content (around 0.05) while the highest was recorded for the fruit weight (0.34). The remaining seven traits ranged from 0.10 to 0.16 (Table 2). This indicated that the non-additive gene effects were more prevalent for the characters under investigation. In general, the GCA variance was higher than that of SCA variance for the characters studied. The broad-sense heritability values were higher (above 0.9) than the narrow-sense heritability(Table 2), showing that selection between hybrids and varieties will be efficient for the improvement of these traits.

**Table 2.** Analysis of variance for GCA and SCA for the ten descriptors in tomato under leaf curl virus conditions including GCA/SCA ratio, and narrow sense (h^2^) and broad sense heritabilities (H^2^).

| Source of Variation | Genotypes<sup>a</sup> | GCA<sup>a</sup> | SCA<sup>a</sup> | Error | GCA/SCA ratio | $h^2$ | $H^2$ |
| --- | --- | --- | --- | --- | --- | --- | --- |
| DF | 65 | 10 | 55 | 65 |  |  |  |
| Dry matter | 0.38*** | 0.66*** | 0.33*** | 0.11 | 0.16 | 0.22 | 0.95 |
| Equatorial diameter | 0.68*** | 0.87*** | 0.65*** | 0.01 | 0.10 | 0.17 | 0.97 |
| Fruit weight | 964.97*** | 2755.90*** | 639.35*** | 31.90 | 0.34 | 0.38 | 0.94 |
| Locules | 0.87*** | 1.48*** | 0.76*** | 0.02 | 0.15 | 0.22 | 0.96 |
| Lycopene | 1.58*** | 1.91*** | 1.51*** | 0.01 | 0.10 | 0.16 | 0.95 |
| Marketable yield | 0.55*** | 0.71*** | 0.52*** | 0.01 | 0.10 | 0.17 | 0.96 |
| Pericarp thickness | 0.03*** | 0.05*** | 0.03*** | 0.07 | 0.13 | 0.20 | 0.95 |
| Polar diameter | 0.83*** | 0.55*** | 0.88*** | 0.01 | 0.05 | 0.10 | 0.97 |
| Total yield | 0.56*** | 0.91*** | 0.50*** | 0.01 | 0.15 | 0.22 | 0.96 |
| TSS | 0.35*** | 0.28*** | 0.36*** | 0.01 | 0.06 | 0.10 | 0.94 |

<sup>a</sup>\*\*\*, \*\*, \* indicate significant at $p < 0.001$ , $p < 0.01$ , or $p < 0.05$ , respectively.

A higher magnitude of additive gene effects is useful for the development of pure-lines and to proceed with selection based breeding approaches. Non-additive gene effects are used for the development of hybrids. In previous studies presence of both additive and non-additive gene action was reported for most of the characters studied in a tomato. For example, additive gene action was reported to control yield and its component traits in tomato[49,50]. In contrast, nonadditive gene action was reported to control many traits in tomato [51,52]. Similarly, the additive and non-additive inheritance of biochemical traits like dry matter, lycopene, and TSS was also reported [53,54].

### 3.3. Bayesian Model-Based Estimates and Predictions of Total Yield

Both GCA and SCA contributed for the total yield showing that both additive and nonadditive effects were significant. However, GCA values for the total yield was higher than the SCA values (Table 2). The predicted means further revealed this with the BayesDiallel Bab model (Figure 2.).

**Figure 2.**
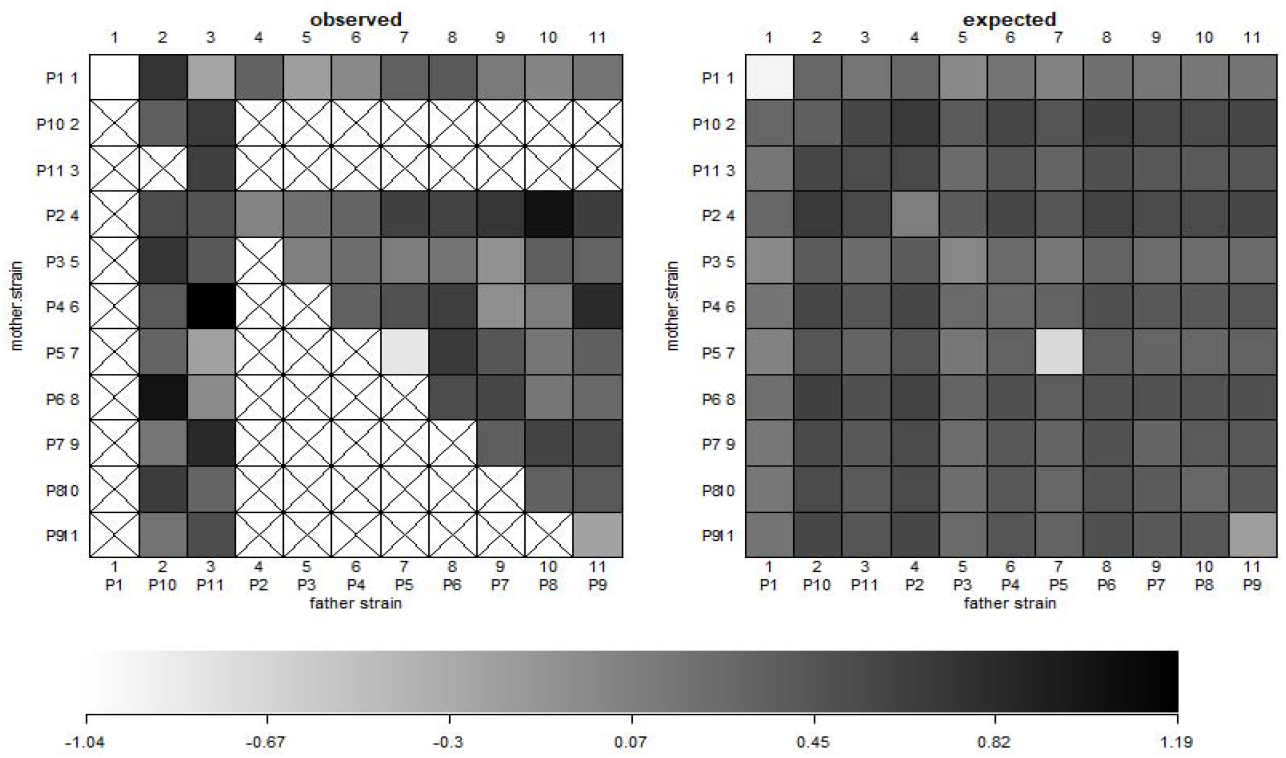
Total yield data of 55 hybrids and 11 parental genotypes in a half diallel. On the left, of parental (P1 to P11), with crossed boxes indicating the missing data (as half diallel was used) and the shaded box representing the values based on the horizontal scale (below). The right side graph shows predictive means based on the Bab diallel model on the scale of (−1.04 to 1.19).

The fixed and individual strain additive effects were more stable and less desperessed, than the parent of origin and inbreeding effect (Figure 3). Justifying a large amount of GCA component identified. The parental genotypes P1, P5 and P9 showed a negative parent of origin and inbreeding effects (Figure 3). The observed vs expected values were found different for some hybrid combinations (Figure 2). It could be attributed to the fact that whiteflies have a preference for some genotypes. The virulence affected the degree and plant reaction to the disease contributed to such effects [55,56].

**Figure 3.**
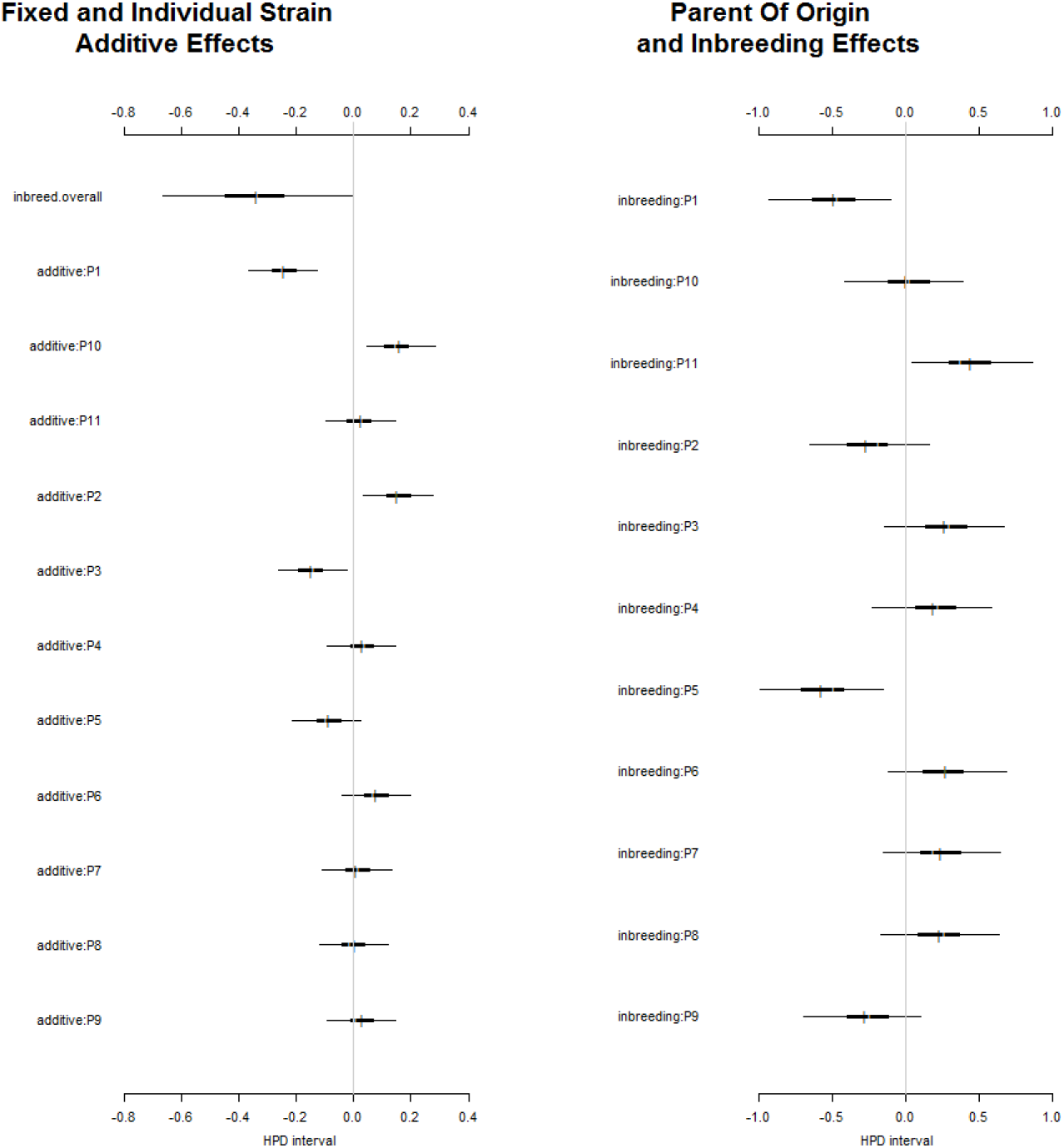
Highest posterior density (HDP) intervals of parent-specific additive effects, parent of origin and inbreeding effect of the 11 parents studied.

Likewise, under TLCV disease pressure Bayesian approach helped in the determination of heritable and non-heritable components influencing the total yield. This precise determination can be used to determine the most appropriate breeding strategy for maximising the genetic gain. Overall, this variance projection approach (VarP) is more precise in providing the information about inheritance classes those will affect the future experiments if these eleven parental genotypes are used again in the future [57]. The total yield included the additive effect (VarP[a]=0.24) while, the effect of non-additive variance was in the form of inbred penalty (VarP[B]=0.07), and the noise was 58.97 for the total yield (Figure 4). Previously, in case of cross-pollinated carrots, it was found that the influence of nonadditive variation was largely due to the overall inbred penalty (non-additive effects), which contributed significantly to canopy height, shoot biomass, and root biomass [35].

**Figure 4.**
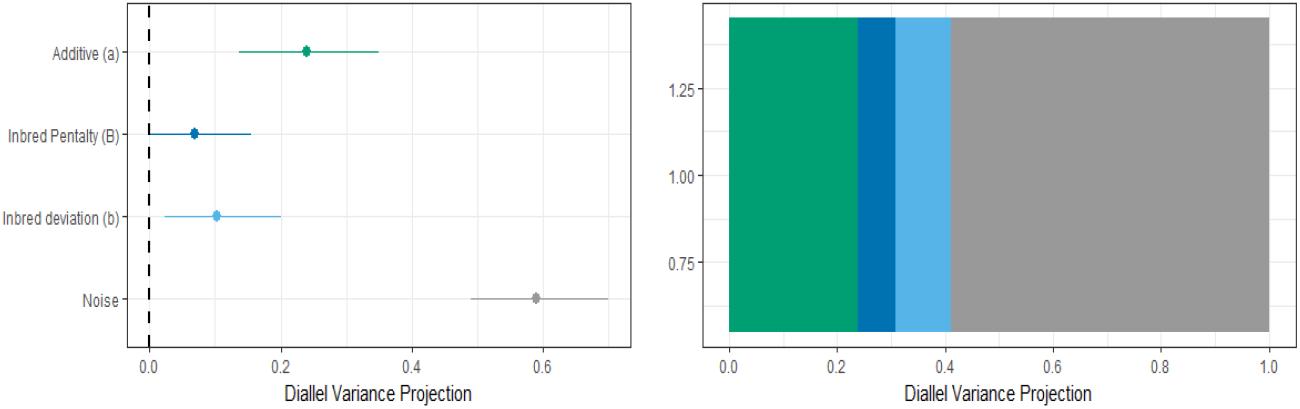
Diallel variance projections the genetic architecture for total yield.

### 3.4. GCA and SCA estimates

Estimates of general combining ability are presented in Table 4. The estimates were highly significant for all the characters studied for dry matter content. P2 expressed the highest GCA effect (0.346. For equatorial diameter, P4 was the best general combiner with the estimates of 0.273. Regarding fruit weight P5 (9.596) was the best general combiner followed by P8 (8.453) and P7 (7.770) (Table 4). The genotype with the highest CGA effects for the number of locules is P2 (0.355) followed by P8 (0.350) and P4 (0.261). The genotypes with above average GCA effects for lycopene content included P3 (0.454), P6 (0.427) and P9 (0.213) (Table 4). The highest GCA effects for marketable yield of 0.188 were recorded in genotypes P2, followed by P11 and P10. For pericarp thickness P8 (0.115) has the highest GCA effects. For polar diameter, highest GCA effects were observed in P7 (0.263). While for total yield P10 (0.215), P11 (0.164) and P6 (0.154) high GCA effects. Likewise, the GCA effects of 0.195 and 0.152 were recorded for TSS content in the parents P8 and P1, respectively.

**Table 4.**
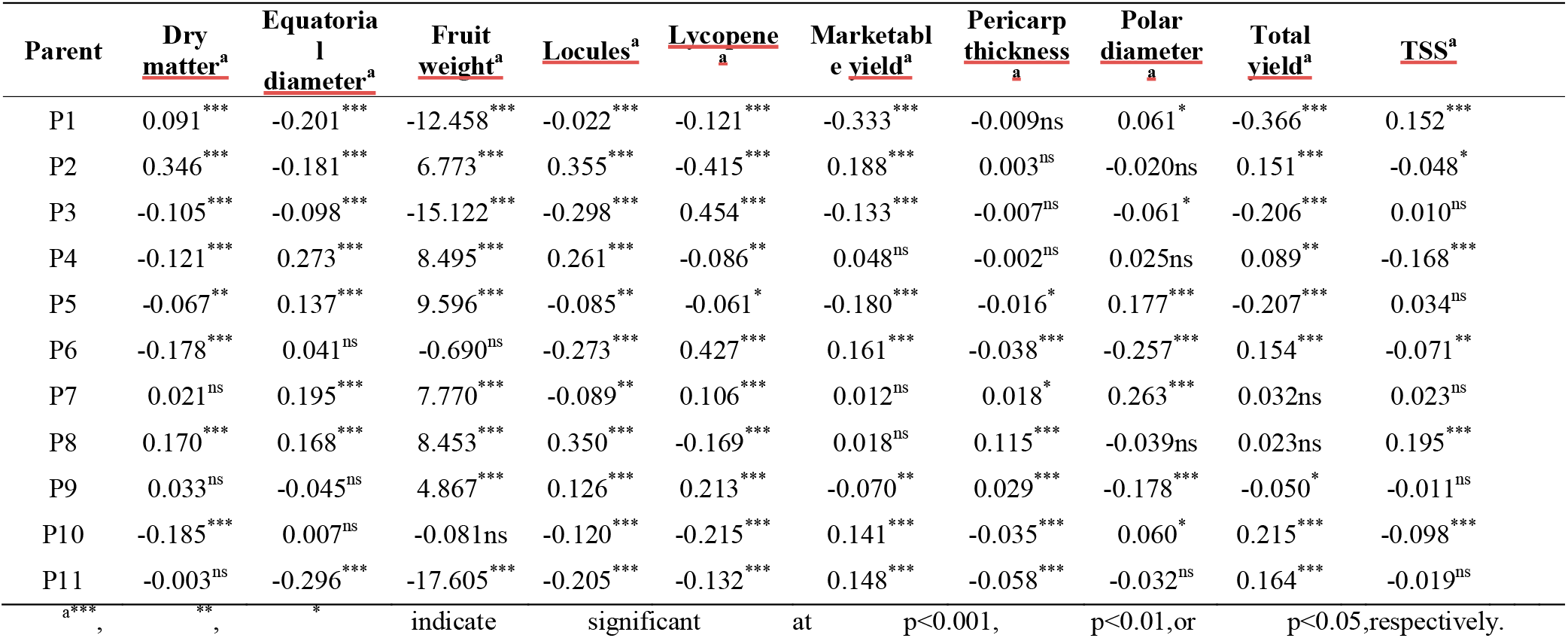
The estimates of general combining ability (GCA) for the parent genotypes (11) for the ten descriptors studied under the influence of leaf curl virus.

The results of specific combining ability estimates are given in Table S1. The crosses with the highest SCA effects for Dry matter content are H22 (1.343), H9 (0.986) and H17 (0.903). The highest SCA effects of 1.386, 1.179 and 1.097 were recorded in the crosses H46, H35 and H23 respectively for the equatorial diameter. For fruit weight, the highest positive SCA effects were observed in crosses H17 (26.728), H35(26.639) and H16 (23.633). For locule number crosses H34, H43 and H46 have the highest SCA effects of 1.551, 1.289 and 1.217 respectively. Similarly, the highest SCA estimates for the lycopene content were recorded in the crosses H30 (1.724), H24 (1.617) and H14 (1.460) in that order. For marketable yield crosses H49 (1.649), H23 (1.134) and H32 (0.918) recorded the highest SCA values. Likewise, for pericarp thickness H29 (0.218), H42 (0.166) and H25 (0.140) were significant. Crosses H3 (1.589), H34 (1.283) and H35 (1.005) possessed the highest SCA values for the polar diameter. In case of total yield H49, H23 and H42 with values of 1.539, 1.157 and 0.917 recorded the highest SCA effects respectively. For the TSS content crosses with highest SCA effects included H49 (1.017), H26 (0.961) and H28 (0.928). Hybrid combinations, H23, H35, H42, H46 and H49 were found to be good specific combiners for more than one trait (Figure 5). However, none of the hybrids exhibited significant SCA effects for all the traits. The information about the SCA and GCA is crucial for maximizing the genetic gain. In case of self-pollinated crops, SCA effects generally do not contribute to the improvement of the trait. These results agree with the results reported in the previous studies under stress and under natural conditions [22,58–60].

**Figure 5.**
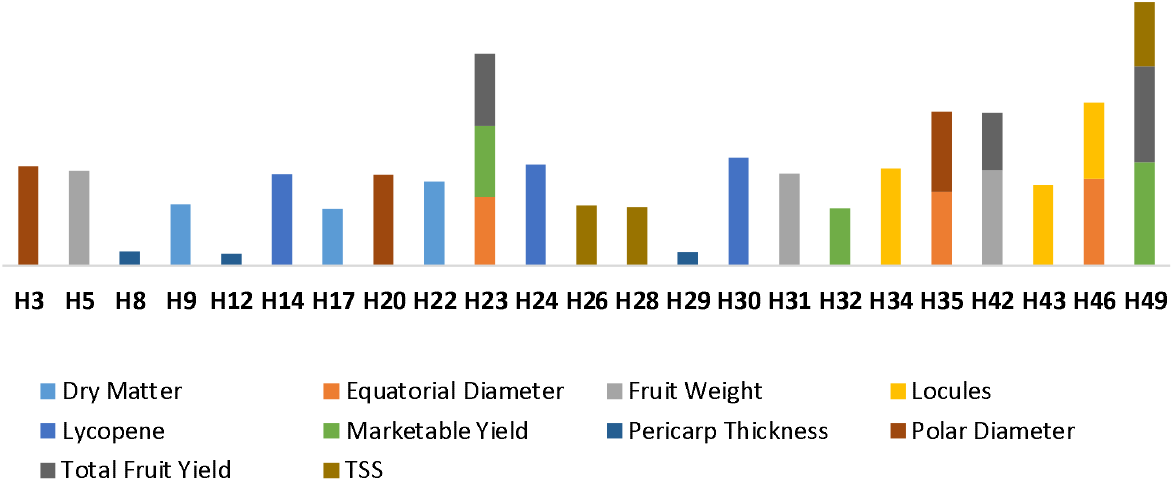
Promising hybrid cross combinations identified based on SCA values, under leaf curl virus infestation.

#### Correlations

A total of 14 correlations were found to be significant (P < 0.05) (Figure 6). One of the correlation was absolute (0.966) that is between total yield and marketable yield. Locule number is found to be correlated with fruit weight (0.43), equatorial diameter (0.32), dry matter (0.31) and TSS (0.17). While the fruit weight was correlated with equatorial diameter (0.54), locules (0.43), pericarp thickness (0.28) and polar diameter (0.27). Previous research works conducted on tomato showed the similar results. In this direction, a high correlation was noticed between fruit yield, fruit weight and pericarp thickness [61,62]. Although it is worth mentioning that no correlation was found between yield and any of the morphological and biochemical traits. Tomato plant yield under TLCV pressure is independent of any morphological and biochemical trait. Likewise, earlier it was shown that under salt stress, plants survival and yield are independent of each other[63].

**Figure 6.**
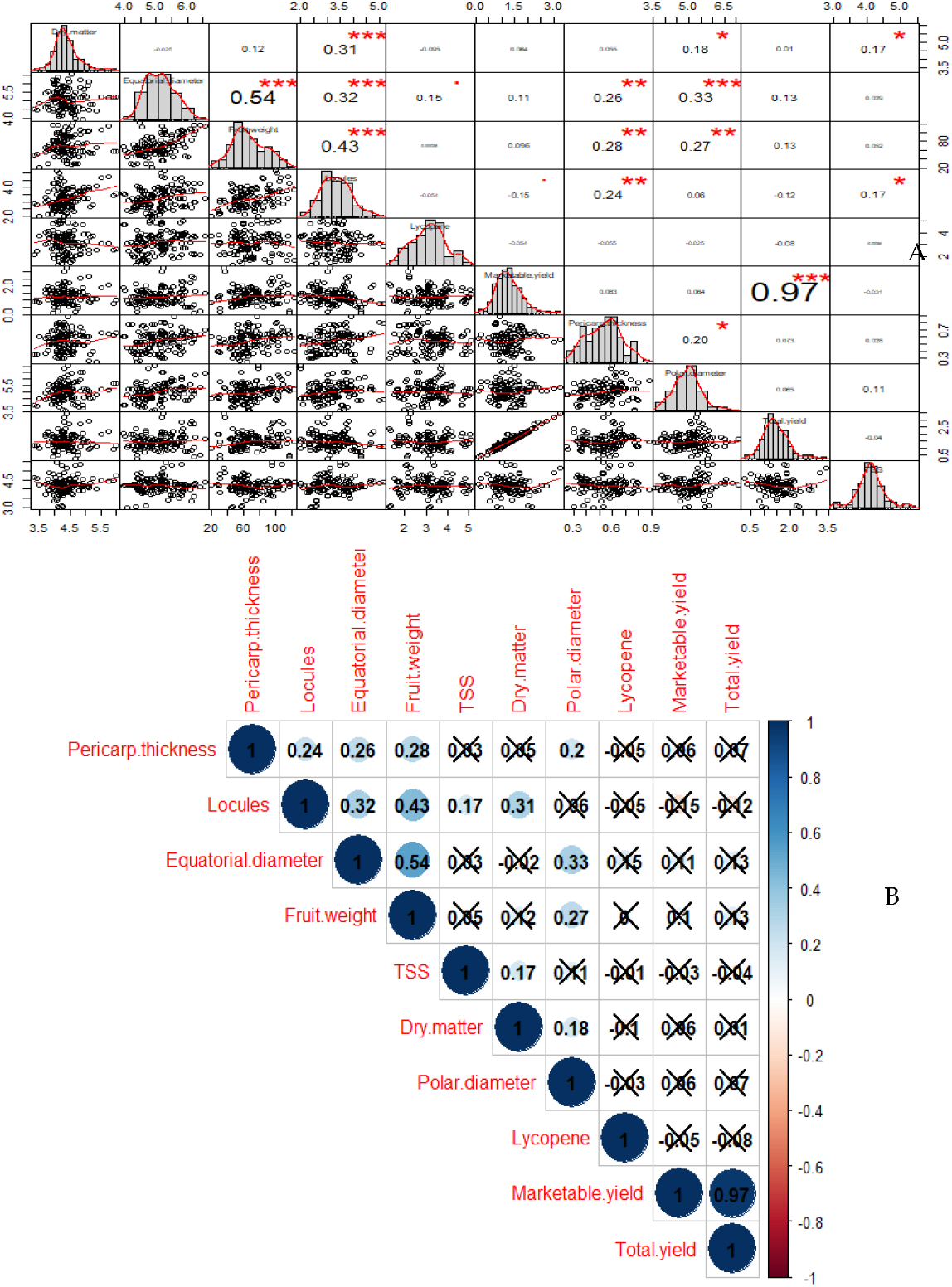
Pearson’s correlation coefficients with significant values at p<0.001 (***), p<0.01 (**), or p<0.05 (*), respectively (upper diagonal) along with the pattern of the distribution of data via scatter plots and histograms (lower diagonal) (A). While all the significant Pearson’s correlation coefficients at p<0.05 (B).

## 4. Conclusions

During the past decades, breeding for tomato leaf curl virus resistance has been a major focus for the resistance breeding programs in tomato. Therefore, breeding efforts have been made to combine significant resistance to TLCV with important fruit quality and yield traits. The diallel matting design is a popular choice as it helps in the identification of parents with good GCA effects and hybrids with good SCA effects. Additionally, it provides the important information on gene action and inheritance of the traits. In this study, we evaluated seven morphological and three biochemical traits of interest for tomato breeding under leaf curl virus pressure. The high diversity in the material was confirmed by GCA and SCA values for all traits. This showed the significance of both the additive and the non-additive effects in the inheritance of the traits evaluated. Also, we have dissected the inheritance of total yield using the Bayesian approach. It was shown that total yield was more dependent on additive variance than the non-additive variance. Overall, this information will be useful to design and develop breeding programs aiming to improve TLCV resistance along with a respectable combination of important traits. The moderately resistant and high yielding parents (P4 and P9) and hybrids (H23,H42, and H49) could be used for the resistance breeding in tomato.

